# MSAGPT: Neural Prompting Protein Structure Prediction via MSA Generative Pre-Training

**DOI:** 10.1101/2024.06.10.598380

**Authors:** Bo Chen, Zhilei Bei, Xingyi Cheng, Pan Li, Jie Tang, Le Song

## Abstract

Multiple Sequence Alignment (MSA) plays a pivotal role in unveiling the evolutionary trajectories of protein families. The accuracy of protein structure predictions is often compromised for protein sequences that lack sufficient homologous information to construct high-quality MSA. Although various methods have been proposed to generate virtual MSA under these conditions, they fall short in comprehensively capturing the intricate co-evolutionary patterns within MSA or require guidance from external oracle models. Here we introduce MSAGPT, a novel approach to prompt protein structure predictions via MSA generative pre-training in the low-MSA regime. MSAGPT employs a simple yet effective 2D evolutionary positional encoding scheme to model the complex evolutionary patterns. Endowed by this, its flexible 1D MSA decoding framework facilitates zero-or few-shot learning. Moreover, we demonstrate that leveraging the feedback from AlphaFold2 can further enhance the model’s capacity via Rejective Fine-tuning (RFT) and Reinforcement Learning from AF2 Feedback (RLAF). Extensive experiments confirm the efficacy of MSAGPT in generating faithful virtual MSA to enhance the structure prediction accuracy (up to +8.5% TM-Score on few-shot scenarios). The transfer learning capabilities also highlight its great potential for facilitating other protein tasks.

## 1 Introduction

The advent of deep learning has significantly propelled progress across various scientific domains, exemplified by breakthroughs such as AlphaFold series [1, 2] for accurate biomolecular interaction predictions, AlphaGeometry [3] for intricate geometry and mathematical reasoning——to name a few. Among these, AlphaFold2 (AF2) represents a landmark within structural biology, achieving an *in silico* precision of approximately 90% atomic accuracy that rivals wet lab experiments on protein structure predictions (PSP). The remarkable success of AF2 can be attributed to its innovative use of co-evolutionary information supported by the Multiple Sequence Alignment (MSA). MSA aggregates homologous sequences from vast databases, providing a comprehensive overview of evolutionary trajectories, which is critical for accurately predicting protein structures [1, 2, 4]. An illustrative example in Figure 1(a) showcases that the correlations analysis among amino acids (AAs) sites could reveal contacts or conservative regions in the folding structure. Unfortunately, not all proteins possess a rich set of homologous counterparts. Statistical investigations reveal that approximately 20% of metagenomic proteins [5] and around 11% of proteins from eukaryotic and viral origins [6] are classified as “orphan” proteins. This presents a significant challenge for MSA-search algorithms in constructing high-quality MSA, consequently impeding the performance of PSP models [2].

**Figure 1:**
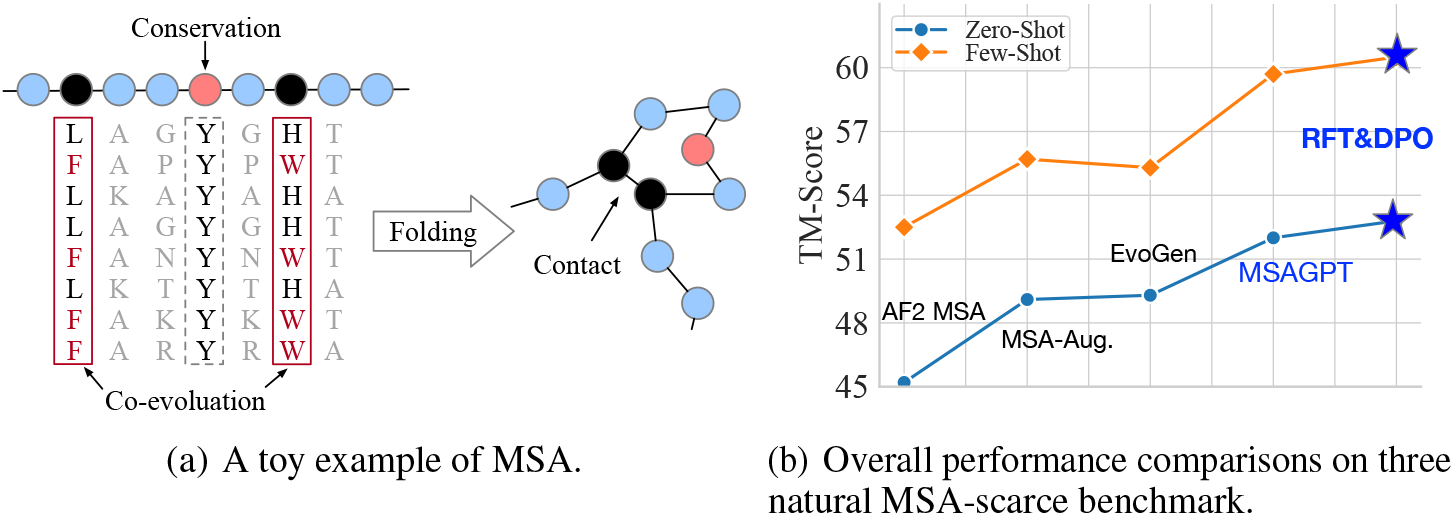
(a) The illustration of MSA and (b) performance comparisons between MSAGPT and advanced baselines.

Drawing on the impressive capabilities of large language models endowed either by the autoencoding [7] or the autoregressive language modeling regime [8, 9], protein language models (PLMs) have been developed to unveil the evolutionary patterns and sequence characteristics intrinsic to protein structures. Specifically, generative PLMs [10, 11, 12], trained on vast protein databases [13, 14, 15, 16] have achieved unparalleled success in generating novel proteins with desired structural properties. These achievements underscore the efficacy of language models in identifying evolutionary patterns within individual protein sequences. Inspired by this, subsequent works [17, 18] attempt to further integrate MSA as the input or by directly generating virtual yet informative MSA [19, 20, 21] to provide additional evolutionary insights. These approaches usually adopt customized attentions that merely allow attention aggregated among specific directions, such as axial attention [22], for separately analyzing the row- and column-wise co-evolutionary patterns in MSA. However, these attention mechanisms usually have low efficiency in capturing the evolutionary information in MSA, or even fail to adequately capture intricate co-evolutionary dynamics. Taking Figure 1(a) as an example, it is imperative to concurrently analyze the pairwise or high-order relationships of amino acid sites across all homologs to deduce the structural constraints influencing the folding structures, which may not achieved by customized attention. The limited capacity may result in compromised performance on the task that highly resorts to co-evolutionary information.

Built upon the insights mentioned above, we introduce MSAGPT, a simple yet effective framework that prompts protein structure prediction via MSA generative pre-training. This method facilitates *de novo* MSA generation, aiding in protein structure prediction in scenarios with limited MSA available. MSAGPT is characterized by its unique features:

- **2D Evolutionary Positional Encoding**. We employ an innovative dual-axis positional encoding scheme that captures column- and row-wise co-evolutionary information concurrently. This method provides a comprehensive understanding of complex evolutionary relationships with high efficacy. enhancing the model’s generative capabilities.
- **1D Zero-/Few-Shot MSA Decoding**. With 2D positional encoding, MSAGPT re-formalizes MSA generation as a one-dimensional sequence generation task, optimized by the simple next-token-prediction objective. This enables MSAGPT to conduct zero- or few-shot MSA generation under a flexible in-context learning framework.
- **Learning from AlphaFold2 Feedback**. MSAGPT further utilizes feedback from AlphaFold2 to reduce hallucinations during MSA generation. This approach ensures the generation of reliable and informative MSA, thus enhancing protein structure prediction.

Extensive experiments conducted on three benchmarks, CAMEO [23], CASP, and PDB [14], demonstrate the superior capacity of MSAGPT in generating high-quality MSA. Notably, MSAGPT outperforms existing MSA generation models on both zero- and few-shot scenarios. Impressively, AF2 with virtual MSA generated by MSAGPT significantly improves the structure prediction accuracy than that with natural MSA on cases with limited homologous information. Moreover, the subsequent Rejective Fine-tuning (RFT) and learning from AF2 feedback (RLAF) stage further enhance the model’s ability to generate informative and faithful MSA, outperforming the original MSAGPT by a large margin, as shown in Figure 1(b). Additionally, we demonstrate that virtual MSA can also benefit other tasks. We expect MSAGPT to become integral in supplementing protein-related tasks requiring critical evolutionary information from MSA. The code, data, and scripts are available https://github.com/THUDM/MSAGPT.

## 2 Related work

### Protein Structure Prediction

Proteins are fundamental to the various biological processes that sustain, grow, and protect living organisms. Groundbreaking deep learning approaches, such as AlphaFold series [1, 2] and RoseTTAFold [4], have been developed to predict the folding structures based on their sequences. These methods have achieved comparable structure prediction accuracy to conventional wet-lab experiments. The success of these cutting-edge methods largely rely on the utilization of MSA, which are retrieved through search algorithms [24, 25, 26, 27] against vast databases [13, 14, 15, 16]. However, challenges arise with “orphan” protein sequences, which lack sufficient homologous sequences for accurate structure prediction. Single-sequence PSP methods [28, 11, 29, 30] are designed to infer folding structures directly from the query protein sequences. Despite these advancements, the accuracy of predictions from single-sequence methodologies generally falls short of those derived from MSA-based algorithms.

### Protein Language Models

Protein Language Models (PLMs) have emerged as a groundbreaking development in computational biology. A family of models including ESM [28, 31], etc [32], are trained on single sequences, towards understanding protein structural features. MSA Transformer [17] further incorporates MSA as the input, achieving better performance than these single sequence models, underscoring the importance of utilizing the evolutionary information from MSA [33, 34, 35]. As for protein design, PLMs including ProGen [10, 36] and ProtGPT2 [12], endowed by the autoregressive training regime, enable the generation of diverse and realistic protein sequences. To enhance MSA generation, MSA-Augmentor [20] and PoET [19] employ the seqs2seqs pretraining, which adopts the sequential axial attention mechanism to capture the evolutionary data across and within the input sequences, both horizontally and vertically. EvoGen [21], serving as the meta generative model, aims at producing virtual MSA for enhancing protein structure predictions. However, it highly resorts to external structural prediction models to optimize its performance.

### 2.1 Aligning with Human Preferences

Aligning language models with human preferences has been shown to be effective in improving generation quality [8, 37, 38, 39]. Learning from human feedback based on pre-trained models is a common approach in achieving this alignment. Existing methods typically employ supervised fine-tuning using human-annotated datasets or reinforcement learning from human feedback pipelines. Reinforcement algorithms such as Proximal Preference Optimization [38] (PPO) and Direct Preference Optimization [37] (DPO) are commonly used in these pipelines. Inspired by these, we utilize the feedback from AlphaFold2 to further enhance the generation capability of the pre-trained model, which helps mitigate hallucinations and enables the model to generate accurate and reliable MSA.

## 3 Preliminary

### Definition 1

***Multiple Sequence Alignment (MSA)*** *Given the query protein sequence Q* ∈ 𝔸^1*×L*^, *where* 𝔸 *denotes the set of alphabetic symbols used to represent the 20 basic amino acids and L represents the number of amino acids per sequence, the MSA M* ∈ 𝔸^*N×L*^ *of Q is comprised of N homogeneous protein sequences, which can be obtained either by searching over protein databases or generating with MSA generation methods*.

### Problem 1

***Prompting Protein Structure Prediction by MSA Generation*** *Given Q with initial MSA M*_*init*_ ∈ 𝔸^*n×L*^ *as the prompt, where n* = 0 *indicates the zero-shot MSA generation and n >* 0 *signifies the few-shot MSA generation, we target at learning a function f to generate virtual MSA M*_*gen*_ ∈ 𝔸^*m×L*^ *based on Q and M*_*init*_, *such that the structure prediction accuracy based on the augmented MSA M*_*aug*_ ∈ 𝔸^(*n*+*m*)*×L*^ *significantly surpasses that based on the initial MSA M*_*init*_,

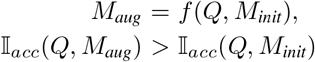

*where the* 𝕀_*acc*_ *is prediction accuracy comparing the prediction result of AF2 and the ground truth*.

In this paper, we mainly focus on improving the structure prediction accuracy in the low-MSA regime, i.e., the cases that lack a sufficient number of homologous sequences.

## 4 Methodology

Given a query sequence and its retrieved natural MSA, we aim to comprehensively understand the co-evolutionary patterns in MSA, such that we can generate informative virtual MSA for prompting protein structure prediction in the low-MSA regime. Conceptually, the co-evolutionary information is analogous to the covariance matrix in mathematics, depicting the correlations among amino acids by comparing pairwise or high-order correlations among amino acid sites in MSA, as depicted in Figure 1(a). To achieve this goal, MSAGPT contains two key adoptions, distinguishing it from existing MSA-based PLMs that rely on customized attentions [2, 17, 20, 19]: **2D Evolutionary Positional Encoding**. Introduces an adaptive dual-axis positional encoding scheme that captures column- and row-wise co-evolutionary information concurrently. And **1D Zero-/Few-Shot MSA Decoding**. Re-formalizes MSA generation as a one-dimensional sequence generation task based on the proposed 2D positional encoding scheme, which enables MSAGPT to conduct zero-or few-shot context learning MSA generation framework. The overall framework is illustrated in Figure 2.

**Figure 2:**
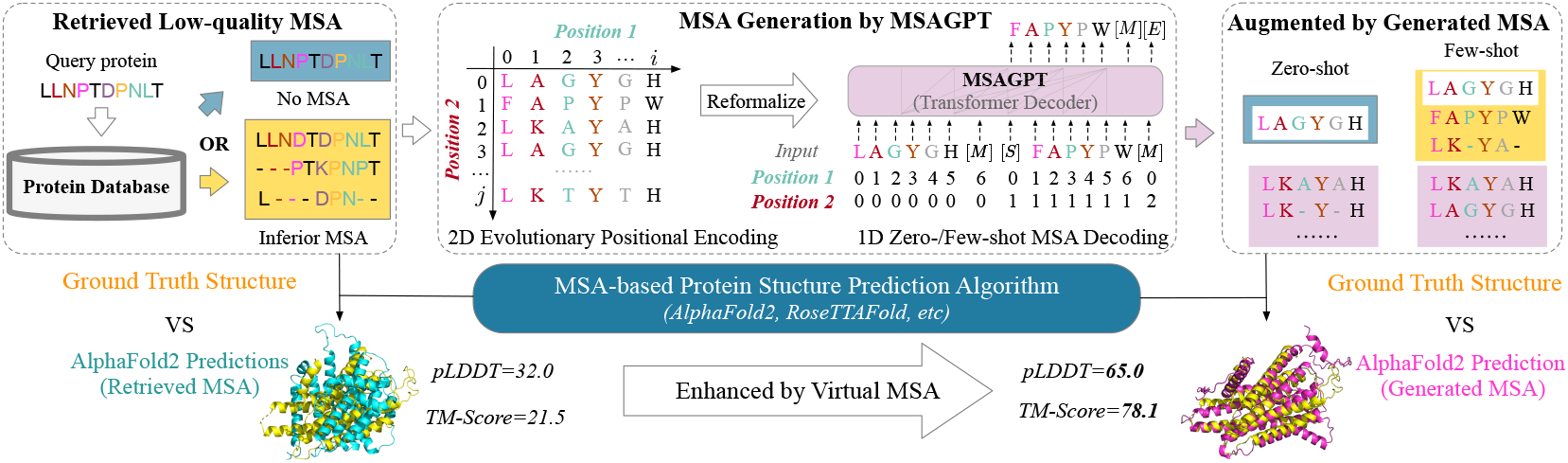
The overall framework of prompting protein structure predictions via MSA generation. **Left**: The challenge faced by conventional search algorithms on protein with scarce homologous sequences, resulting in suboptimal alignments. **Middle-to-Right**: MSAGPT generates informative and high-quality MSA for such challenging queries, presenting a promising approach to overcoming these limitations. `[M]` denotes the sequence separator. `[S]`, `[E]` are the special tokens to represent the start or end of MSA generation.

### 4.1 2D Evolutionary Positional Encoding

Vanilla transformers typically use 1D positional embeddings to incorporate absolute and relative positional information of tokens. However, when dealing with MSA, which represents stacked homologs, the structure is different. Each row of MSA corresponds to a distinct protein sequence, while each column represents the evolutionary trajectories of a specific amino acids (AAs) site. To effectively capture the evolutionary patterns, recent approaches [2, 17, 20] have employed decoupled axial attentions, which are designed to capture explicit co-evolutionary information along the rows (protein sequences) and columns (AAs sites). However, these methods often suffer from low efficiency in capturing the information dynamics or fail to capture the evolutionary information adequately.

In light of this, we introduce a novel two-dimensional evolutionary positional encoding scheme, illustrated in Figure 2. Given an MSA **M** ∈ 𝔸^*N×L*^, we define a 2D positional id matrix **P** ∈ ℕ^2*×N×L*^, where the first positional id **P**_**0**_ ∈ ℕ^1*×N×L*^ indicates the column position, i.e., *P*_0_[*i*,·] = {0, 1, · · ·, *L*}, and the second positional id **P**_**1**_ indicates the row position, i.e.,*P*_1_[*j*, ·] = {0, 1, · · ·, *N*}. The two positional ids are projected into two vectors added to the input token embeddings. We utilize the Rotary Positional Encoding (RoPE) [40] technique, specifically adapting its two-dimensional variant^4^ to suit our 2D positional encoding requirements.

#### Comparison with Axial Attentions

Considering the 2D positional id (*P*_0_, *P*_1_), the self-attention among AAs (*α, β*) meets the following unit patterns, as illustrated in Figure 3:

**Figure 3:**
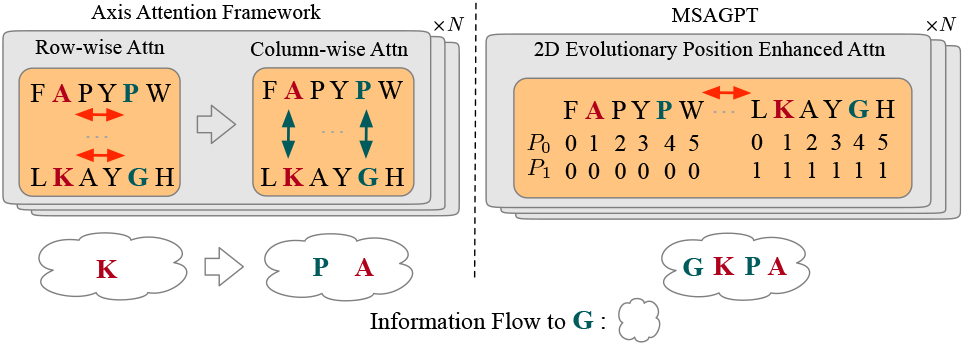
Comparisons among the axial attention (exemplified by [17]) and the one in MSAGPT in a single layer. Here we focus on the information aggregated to the AA “G”. The 2D evolutionary position enhanced attention shows higher efficiency than the decoupled axial attentions with one-step aggregation to attain sufficient information.

- 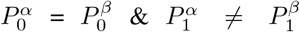. Indicates *α* and *β* reside in the same site across different protein sequences, such as the AA pair (A, K) and (P, G), enabling column-wise self-attention that highlights evolutionary patterns conserved across sequences.
- 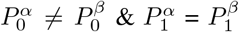. Suggests *α* and *β* are positioned in the same protein sequence but at different sites, such as the AA pair (A, P) and (K, G), facilitating row-wise self-attention that captures sequence-specific features.
- 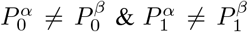. Denotes *α*and *β* lack explicit correlation, such as the AA pair (A, G) and (P, K), may be serving as anchor nodes for complex coevolutionary information diffusion.

Conceptually, the 2D positional encoding encapsulates the explicit row- and column-wise self-attention patterns with high efficacy. Moreover, it allows unrestricted information diffusion, that is, enabling any two amino acids to attend to one another. Such a framework facilitates unveiling complex co-evolutionary patterns, such as high-order correlations among AAs, that customized self-attentions might overlook.

### 4.2 1D Zero-/Few-Shot MSA Decoding

Leveraging the 2D evolutionary positional encoding, we further release the stacked MSA decoding task into the scalable 1D sequence generation framework, without compromising the integrity of co-evolutionary information. Specifically, we convert the MSA **M** ∈ 𝔸^*N×L*^ into the flatted 1D sequence **M**^*f*^ ∈ 𝔸^1*×NL*^. Similarly, the 2D positional id matrix **P** ∈ ℕ^2*×N×L*^ is reshaped into a flattened format, **P**^*f*^ ∈ ℕ^1*×*2*×NL*^. This allows the model to conduct a simple auto-regressive generation process, as illustrated in Figure 2.

## Discussions

Admittedly, introducing 2D positional encoding introduces higher time complexity in comparison to conventional customized attention mechanisms (from *O*(*N* ^2^*L*) + *O*(*NL*^2^) to *O*(*N* ^2^*L*^2^)). However, it is worth noting that the original stacked nature of MSA poses challenges for integrating it with acceleration techniques used in large language models, such as Flash Attention [41,42]. The 1D decoding framework, conversely, can be easily scaled to accommodate in-context learning frameworks, such as retrieval augmented generation, to further enhance the model’s generation capability and expand its application range. From a practical standpoint, the high parallelism of the 1D decoding framework demonstrates superior inference speed, benefiting from techniques like Flash Attention and KV-cache, while incurring negligible memory overhead compared to customized attention mechanisms. For further details, please refer to Appendix Section A.4.

## 5 The Training Pipeline of MSAGPT

The training pipeline involves three successive stages: **Stage 1: MSA Generative Pre-Training** to obtain the base MSA generation model; **Stage 2: Rejective Fine-tuning (RFT)** to instruct the base model with high-quality MSAs via AF2 annotations, which can reduce generation hallucinations ; **Stage 3: Reinforcement Learning from AlphaFold2 Feedback (RLAF)** to further enhance RFT model’s capabilities based on the feedback of AF2. (See Appendix Section A for training details.)

### 5.1 Stage 1: MSA Generative Pre-Training

#### Pre-Training Dataset

We utilize the Uniclust30 MSA dataset from OpenProteinSet [43], which is processed through an all-against-all search on Uniclust30 [44] using HHblits [45]. This results in approximately 16 million MSAs

#### Pre-training Objective

We adapt the language modeling objective [46] to the MSA generation task. The cross-entropy loss for modeling the intrinsic distribution of MSA **M**^*f*^ ∈ 𝔸^1*×NL*^ is defined as:

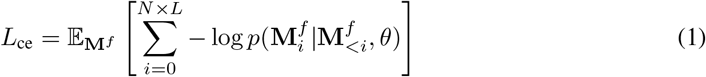

where **M**^*f*^ ∈ 𝔸^1*×NL*^ is 1D flatted version of the input MSA and *θ* is the learned parameter.

### 5.2 Stage 2: Rejective Fine-tuning (RFT)

Noted that the pre-trained dataset inevitably contains noisy co-evolutionary patterns, such as large portions of deletions and insertions, which may mislead the base model to yield hallucinated cases, i.e., the linguistically reasonable but intrinsically unfaithful MSA. Thus we select highly-quality MSAs to further fine-tune the base model via a rejective sampling procedure based on the AF2-annotation.

#### RFT Dataset

We collect 120,780 protein sequences with structures from Protein Data Bank (PDB) [14]. For the sequence *Q*, we search its MSA *M* ∈ 𝔸^*N×L*^ from UniClust30 [44] with HHblits [45]. Then we sample several MSA subsets **m** = {*m*_1_, *m*_2_, …, *m*_*i*_} with replacement, where *m*_*i*_ ∈ 𝔸^*n×L*^ and *n* ≪ *N*. To assure the information density of the sampled data, we filter out the MSA with depth *N* fewer than ⌈*n* × *i/*2⌉. Subsequently, we employ AF2 to score the sampled subset using the structure prediction accuracy 𝕀_*acc*_(*Q, m*_*i*_). Then the RFT dataset 𝒟_RFT_ is defined as:

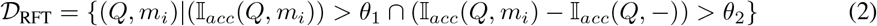

where 𝕀_*acc*_(*Q*, −) indicates the prediction accuracy without using MSAs. Practically, we set the sampling number *i* = 10, the depth of each sampled MSA subset *n* = 16, *θ*_1_ = 0.9, and *θ*_2_ = 0.2, which results in 𝒟_RFT_ of approximately 60k samples. The base model is fine-tuned on 𝒟_RFT_ with the same pre-training objective.

### 5.3 Stage 3: Reinforcement Learning from AlphaFold2 Feedback (RLAF)

We further employ AF2 as the reward model to perform the Reinforcement Learning with AF2 Feedback (RLAF) using Direct Preference Optimization [37] (DPO) to further guide the RFT model to decode meaningful structure-related MSA patterns that align with the preference of AF2.

#### RLAF Preference Dataset

For each query *Q* from the PDB, we use the RFT model to generate its MSA *M* ∈ 𝔸^*N×L*^ in zero-shot manner. Then, we also sample several MSA subsets **m** = {*m*, *m*, …, *m* } and obtain the preference dataset 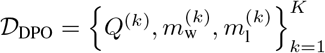 as follows,

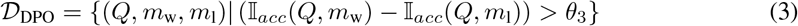

where we set the *θ*_3_ = 0.3, rendering the number of preference data *D*_DPO_ = 11*k*.

#### RLAF Training Objective

The adapted DPO loss is defined as:

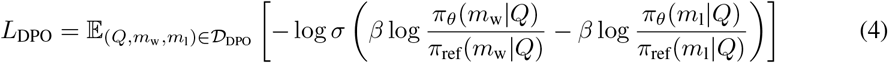

where *π*_*θ*_ and *π*_ref_ are initialized by the RFT model and *π*_ref_ is frozen while *π*_*θ*_ is optimized. During the RLAF training phase, we found that merely using the DPO loss led to training instability. Thus we also adopt the pre-training loss *L*_ce_ for the chosen answer *m*_*w*_ as a regularization term with the coefficient factor *λ* in the total loss to mitigate this issue. The total loss *L* = *L*_DPO_ + *λL*_CE_, *λ* = 0.1. Another critical coefficient *β*, which measures the penalty intensity of DPO for incorrect answers is set to *β* = 0.1.

## 6 Experiments

### 6.1 Setup

#### Benchmarked Dataset

We employ the datasets from CAMEO [23], CASP14&15, and PDB [14], which are esteemed benchmarks in protein structure analysis spanning a diverse array of biological protein families. For each protein sequence, we search its MSA on UniClust30 database [44] using HHblits [45]. Given our focus on addressing the challenge presented by cases with limited MSA information, we build the benchmark to represent the real-world MSA-scarce conditions. More specifically, we identify 200 protein sequences with the number of searched MSA fewer than 20 (8 from cameo, 13 from CASP14&15, 179 from PDB). All MSA of sequences from the test set are removed from the pre-train dataset (see Appendix Figure 9 for details).

#### Baselines

To assess the performance of MSAGPT, we adopt AF2 as the benchmark MSA-based PSP algorithm. For MSA generation baselines, we compare **MSAGPT**, its RFT-version and its DPO-version with two advanced MSA generation algorithms: **MSA-Augmentor** [20], which utilizes a sequences-to-sequences pre-training architecture incorporating an encoder and a decoder based on the axial attention [17]; and **EvoGen** [21], which employs a meta generative model framework with customized attention, leveraging guidance from AF2 to refine its MSA generation. As PoET [19] is designed for mutational scanning tasks, we don’t take it as the baseline. Additionally, we include the reference model **AF2 MSA**, which utilizes all the searched natural MSA for prediction.

#### MSA Generation Pipeline

Given that MSAGPT can perform flexible zero- or few-shot MSA generation to accommodate different levels of available evolutionary information, we define two generation settings to evaluate models’ performances under varying conditions:

##### Zero-Shot Generation

MSA generation is conducted using only the query sequence as input, emphasizing the model’s ability to infer necessary evolutionary patterns without additional contexts.

##### Few-Shot Generation

All the searched natural MSA are viewed as the prompt to inform the few-shot MSA generation process. Then the generated MSA, combined with the initial prompts, serves as augmented data for structure predictions.

#### Evaluation Metric

We employ TM-Score, a widely-used metric for assessing the structural similarity between predicted structures and ground truth, and pLDDT, a per-residue measure of local confidence, as metrics. All metrics are scaled from 0 to 100 (See Appendix Section B for details).

### 6.2 MSAGPT’s Virtual MSAs Reflect the Co-evolutionary Information

Table 1 showcases the comparative results in three datasets across different baselines. Notably, AF2 MSA, which relies solely on the limited searched MSA without incorporating virtual MSA, exhibits the worst performance. Predictions enhanced with MSA generated by MSA-Augmentor or EvoGen surpass the performance of AF2 MSA. This underscores the critical role of high-quality MSA in enhancing the accuracy of cutting-edge PSP algorithms. Overall, MSAGPT surpasses other advanced baselines by a large margin, achieving +1.4% improvement on CAMEO, +8.5% on CASP, and +4.7% on PDB, as measured by TM-Score. This significant improvement demonstrates not only the superior accuracy and effectiveness of MSAGPT but also its robustness in handling cases with noisy or low-quality MSA. Compared with the base model, the RFT and DPO models achieve higher TM-Score but with lower pLDDT values. This discrepancy might arise from the presence of highly confident (according to pLDDT) but lower-scored decoys (according to TM-Score), as observed in [21], indicating that aligning with the preference dataset, which is filtered based on TM-Score, makes the model more inclined to generate truly informative MSA rather than hallucinated ones. Statistically, MSAGPT effectively improves the prediction accuracy for 91.0% and 88.9% of protein sequences with limited MSA when compared to AF2 MSA on Zero-Shot and Few-shot scenarios, respectively. This significant finding highlights the potential of our MSAGPT framework to uncover and leverage co-evolutionary patterns within bio-sequences.

**Table 1:** The performance of structure prediction on three benchmarked datasets. avg. Depth represents the average depth of searched MSA across all query sequences. Compared with the base model, the RFT and DPO models achieve higher TM-Score while with lower pLDDT values.

| Model | CAMEO<br>(avg. Depth = 8.5) |  |  |  | CASP<br>(avg. Depth = 4.6) |  |  |  | PDB<br>(avg. Depth = 2.6) |  |  |  |
| --- | --- | --- | --- | --- | --- | --- | --- | --- | --- | --- | --- | --- |
|  | Zero-Shot |  | Few-Shot |  | Zero-Shot |  | Few-Shot |  | Zero-Shot |  | Few-Shot |  |
|  | pLDDT | TM | pLDDT | TM | pLDDT | TM | pLDDT | TM | pLDDT | TM | pLDDT | TM |
| AF2 MSA | 63.8 | 55.4 | 77.4 | 71.4 | 44.0 | 32.6 | 54.2 | 44.1 | 55.2 | 45.6 | 61.0 | 52.3 |
| MSA-Aug. | 67.7 | 59.2 | 77.4 | 72.1 | 56.8 | 36.6 | 63.4 | 46.3 | 61.9 | 49.8 | 66.0 | 55.3 |
| EvoGen | 66.1 | 60.3 | 78.6 | 75.3 | 48.2 | 38.4 | 55.1 | 48.5 | 57.6 | 49.5 | 62.8 | 55.4 |
| MSAGPT | **70.8** | 61.4 | **80.8** | 75.2 | **59.0** | 39.8 | **65.4** | 51.0 | **68.6** | 53.4 | **71.3** | 59.6 |
| + RFT | 68.0 | 60.5 | 79.8 | 76.4 | 56.8 | 40.2 | 64.0 | 53.6 | 66.8 | 53.4 | 70.3 | **60.1** |
| + DPO | 68.9 | **62.7** | 80.2 | **76.7** | 54.2 | **43.7** | 62.7 | **57.0** | 64.5 | **53.6** | 68.0 | 59.7 |
|  | (+3.1) | (+2.4) | (+2.2) | (+1.4) | (+2.2) | (+5.3) | (+2.0) | (+8.5) | (+6.7) | (+3.8) | (+5.3) | (+4.7) |

### 6.3 Rethinking the MSA Selection Strategy

We further study the effect of different depths of virtual MSA, as shown in Figure 4(a). We observe a trend where the relative improvement in structure prediction accuracy decreases as the depth of virtual MSA increases. The accuracy based on MSA with 64 MSA sequences even underperforms those based on only 16 or 32 sequences. We hypothesize that increasing the number of virtual MSA beyond a certain threshold may introduce a dilution effect, where the density of valuable co-evolutionary signals is compromised by the inclusion of the hallucinated generation noise. To alleviate this, we explore MSA selection strategies for filtering out low-quality, noise-inducing sequences while retaining those that contribute positively to the accuracy of structure predictions, as illustrated in Figure 4(b) (See Appendix Section C for details).

**Figure 4:**
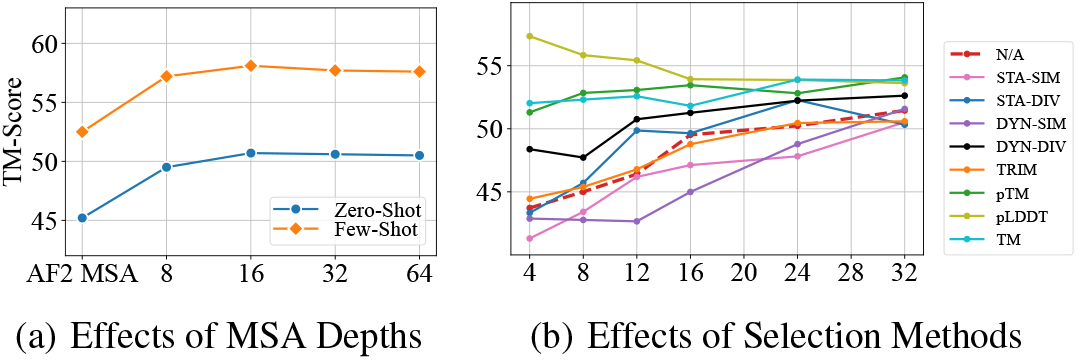
The effect of different MSA depths and selection methods. The X-axis indicates the different MSA depths. The Y-axis represents the TM-Score. The dashed line denotes the non-selection baseline.

#### 1D Sequence Similarity or Diversity Measure

We first arrange MSA by their similarity to the query sequence in descending order. The results reveal that prioritizing MSA based on their high similarity to the query, termed as *static similarity (STA-SIM)*, does not improve prediction accuracy compared to the non-selection approach (N/A). On the contrary, the *static diversity (STA-DIV)* strategy, which favors MSA with lower similarity rankings, slightly outperforms the baseline, highlighting the importance of sequence diversity in enhancing MSA quality. Moreover, we employ the dynamic approach, initially selecting the most (or least) similar MSA to the query sequence and progressively incorporating additional MSA based on their average similarity to the cumulatively selected set, termed as *dynamic similarity (DYN-SIM)* and *dynamic diversity (DYN-DIV)*.

The results further confirm the advantage of fostering diversity within MSA rather than selecting only the sequences with high similarities to the query sequence. We also inspect the effectiveness of the widely-adopted MSA *trimming (TRIM)* strategy [21], which yields a similar TM-Score to the non-selection baseline, undermining its efficacy in selecting MSA with high quality.

#### 3D Structure Affinity Measure

We assume that the generated sequence with high quality should exhibit structural congruity with the query sequence, thereby emitting strong co-evolutionary signals. To validate this, we rank sequences within MSA by their predicted tertiary structures according to the pTM, a predicted TM score [2], pLDDT, and TM-Score, from highest to lowest. These approaches, especially when guided by the pLDDT score, consistently select high-quality MSA, evidenced by the enhanced TM-Score. We compare the non-selection methods (N/A) and pLDDT selection methods on the three benchmarked datasets on few-shot generation scenarios in Table 2. This confirms our hypothesis that structural similarity plays a crucial role in effective MSA selections.

**Table 2:** Performance comparison between non-selection and pLDDT-selection models.

| Model | CAMEO | CASP | PDB |
| --- | --- | --- | --- |
|  | TM | TM | TM |
| MSAGPT-DPO | 76.7 | 57.0 | 59.7 |
| + pLDDT Selection | 77.5 | 57.6 | 60.5 |

### 6.4 Transfer Learning of MSAGPT

Since protein structures largely dictate their functions, the virtual MSA, enhancing structure prediction, should similarly benefit other protein tasks. To validate this, we focus on two protein structural tasks: Contact Prediction (CtP) and Secondary Structural Prediction (SsP) and two protein functional tasks: Localization Prediction (LocP) and Metal Ion Binding (MIB) [11]. We sample 1,000 sequences from each benchmark and conduct 5-fold cross-validation (See Appendix Section B.2 for details).

## Results

Table 3 demonstrate that incorporating MSA from MSAGPT consistently surpasses merely using the single sequence on most tasks. However, it achieves inferior performance on the LocP task, which agrees with the observation [47] that protein language models may not present scaling behavior on several protein functional or property prediction tasks. Nevertheless, the results show the great potential of MSAGPT to contribute to a wider range of protein-related tasks with generated MSA. We are motivated to explore additional transfer tasks to assess MSAGPT’s utility across various domains further.

**Table 3:** Performance comparison between with or without virtual MSA generated by MSAGPT on four protein tasks. ACC is short for Accuracy.

| Model | CtP | SsP | LocP | MIB |
| --- | --- | --- | --- | --- |
|  | ACC | ACC | ACC | ACC |
| w/o Virtual MSA | 11.6 | 66.5 | **58.3** | 57.5 |
| w/ Virtual MSA | **13.1** | **69.0** | 56.4 | **60.3** |

### 6.5 Ablation Study

To understand the effect of various positional encoding strategies on capturing co-evolutionary patterns, we design four model variants: **1D_gpt**: Adopts the standard GPT positional encoding; **1D_2nd**: Utilizes only the second-dimensional of the 2D evolutionary positional encoding mechanism; **1D_1st**: Utilizes the first-dimensional positional encoding; **2D_full**: Implements the 2D evolutionary positional encoding mechanism (See Appendix Section B for details).

## Results

Figure 5 showcases the TM-score distribution across different model variants. The 1D_gpt exhibits the lowest performance, attributed to its simplistic approach of treating the MSA as a concatenation of homologous sequences, thereby failing to discern any co-evolutionary patterns. Both the 1D_1st and 1D_2nd demonstrate significant improvement over 1D_gpt, by explicitly encoding column- or row-wise relationships within the MSA, respectively. Notably, the performance of 1D_1st is better than that of 1D_2nd, suggesting that column-wise covariance patterns play a more crucial role in structural predictions than row-wise patterns. This aligns with the understanding that the permutation of sequence order does not alter the covariance information among residue sites [17]. Remarkably, the 2D_full variant, which incorporates the proposed 2D evolutionary positional encoding, outperforms all other models, which underscores its effectiveness in capturing the intricate evolutionary information present in MSA.

**Figure 5:**
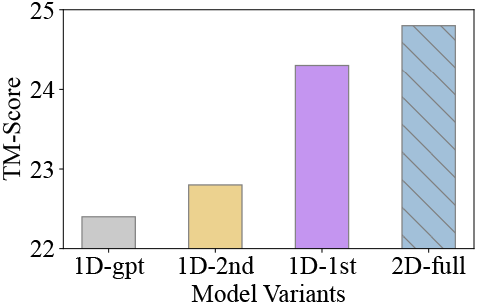
Ablation study with positional encoding variants.

## 7 Limitations

In this section, we discuss some limitations that should be resolved in future work.

### Scaling behavior of MSAGPT

While we have showcased the effectiveness of MSAGPT in generating informative virtual MSA, it is important to note that our pre-training was conducted with a model containing 2.8 billion parameters. The performance and behavior of MSAGPT, when scaled concerning dataset size, model size, and total compute resources, remain unknown.

### General scenarios with sufficient homologs

The primary objective of this paper is to improve the accuracy of protein structure prediction in cases where there is a scarcity of homologous sequences.

However, whether we can further enhance the accuracy in scenarios where there are already sufficient MSA available, augmented by virtual MSA, remains an open question.

### Transfer Learning on a wide range of tasks

While we have demonstrated the transferability of MSAGPT on several tasks, including protein structure prediction and protein function prediction, its performance on a broader range of tasks remains an open question. The ability of a model to transfer its learned knowledge and adapt to new tasks is a critical aspect of transfer learning. While MSAGPT has shown promising results on the tasks it was evaluated on, it is important to assess its performance on a more diverse set of tasks spanning various domains and problem types.

## 8 Border Impact

The aim of this paper is to improve the accuracy of protein structure prediction in cases with limited homologous sequences. The generated MSA also shows great potential to transfer to other protein-related tasks. By leveraging the information encoded in the generated MSAs, it is possible to enhance the performance of various protein-related tasks beyond structure prediction. However, the generative MSA may be misused to contaminate the high-quality nature MSA databases. Thus, it is necessary to train a classifier to distinguish the real and MSAGPT-generated MSA according to the intrinsic features.

## 9 Conclusion

This paper introduces MSAGPT, a novel approach that prompts protein structure prediction via MSA generative pre-training, to enable accurate protein structure predictions in situations where co-evolutionary information is scarce. To meticulously characterize the co-evolutionary patterns within MSA, MSAGPT designs two innovative techniques: the 2D Evolutionary Positional Encoding scheme and the 1D Zero-/Few-Shot MSA Decoding mechanisms. The post-alignment learning from AlphaFold2 feedback further enhances the quality of MSA generation. Empirical experiments conducted on a variety of benchmarks have demonstrated MSAGPT’s robustness and effectiveness. In the future, we plan to apply MSAGPT to broader areas, particularly for tasks that heavily rely on co-evolutionary information, and investigate the aforementioned limitations.

## A Training Settings and Hyper-parameter Studies

The overall training pipeline is illustrated in Figure 6.

**Figure 6:**
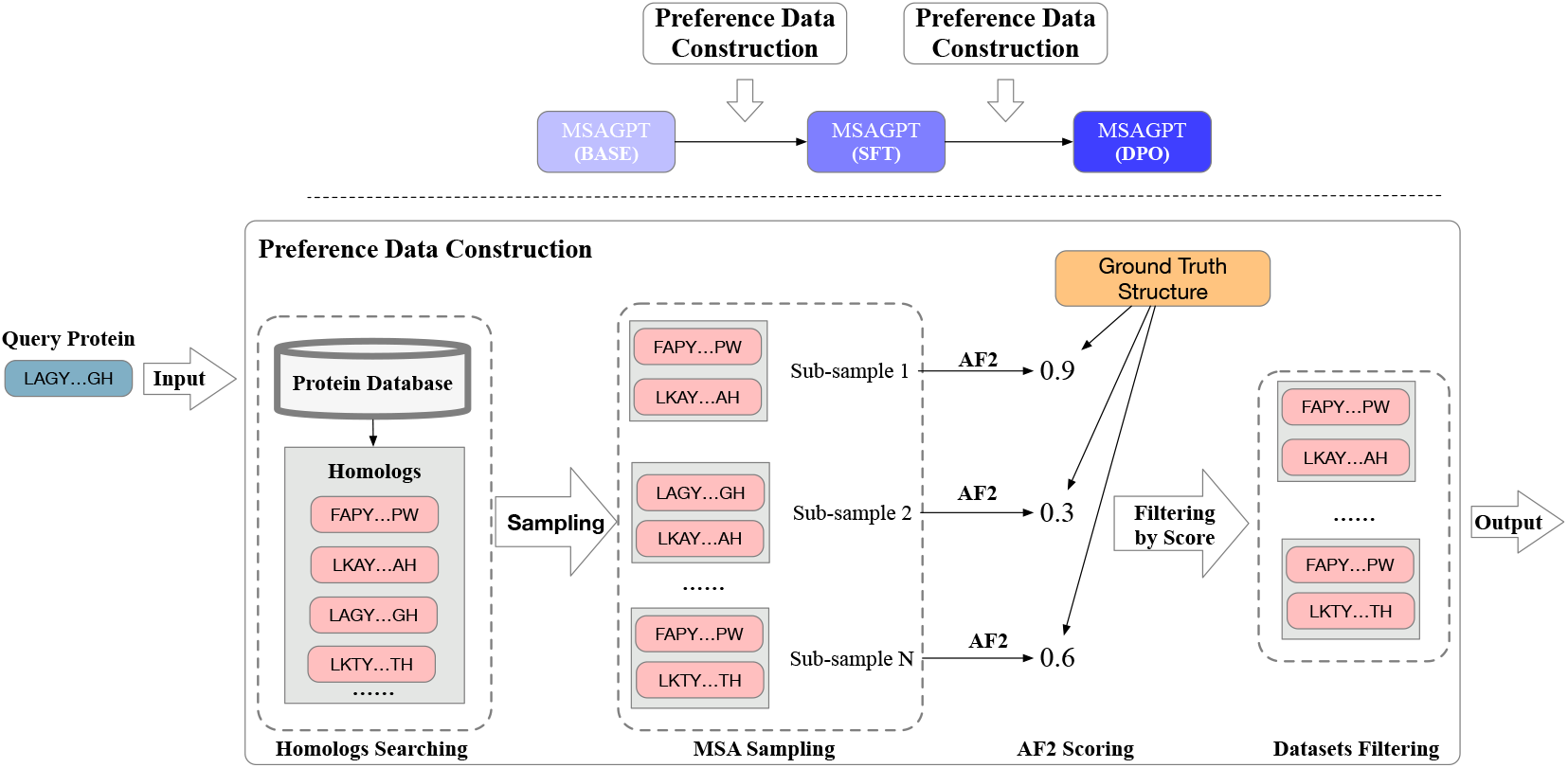
The overall training pipeline and the illustration of preference dataset construction process for SFT and DPO learning stages.

### A.1 Pre-Training

To obtain high-quality MSA, we first screen out clusters with sequence lengths from 25 to 2000, and only retain sequences with the minimum identity of 30% and the largest proportion of gap tokens no more than 10%. The clusters with more than 10 sequences are left. We randomly shuffle the sequences in the MSA to avoid injecting the order bias. Regarding the backbone of MSAGPT, we employ the standard transformer decoder framework [46, 48] and train the model with 2.8 billion parameters owning 36 layers, 2560 embedding size, and 40 attention heads. We employ batches of 48 MSAs with each MSA containing 12,288 residues. We follow BF16 mixed-precision pre-training strategy. We use AdamW [49] as our optimizer with *β*_1_ = 0.9, *β*_2_ = 0.95, eps = 10^−8^ and a learning rate of 1.2× 10^−4^. We use a cosine learning rate schedule, with a warmup of the first 2.5% steps, and decay the final learning rate down to 10% of the peak learning rate. We use a weight decay of 0.1 and gradient clipping of 1.0 without dropout. For the tokenization of the protein data, we use the residue-level tokenizer which is adopted in several PLMs [28, 11, 10]. To save the GPU memory and accelerate the pre-training process, we substitute the standard self-attention module with the Flash Attention-v1 [41] in each layer. All models are trained on 24 A800 GPUs for 254k updates, consuming about 150 billion tokens. This process consumes approximately 150 billion tokens, requiring around 2.7 x 10^18^ floating point operations (FLOPs).

#### Statistics of Pre-trained Dataset

Figure 7 illustrates the length and depth distribution of the pre-training dataset.

**Figure 7:**
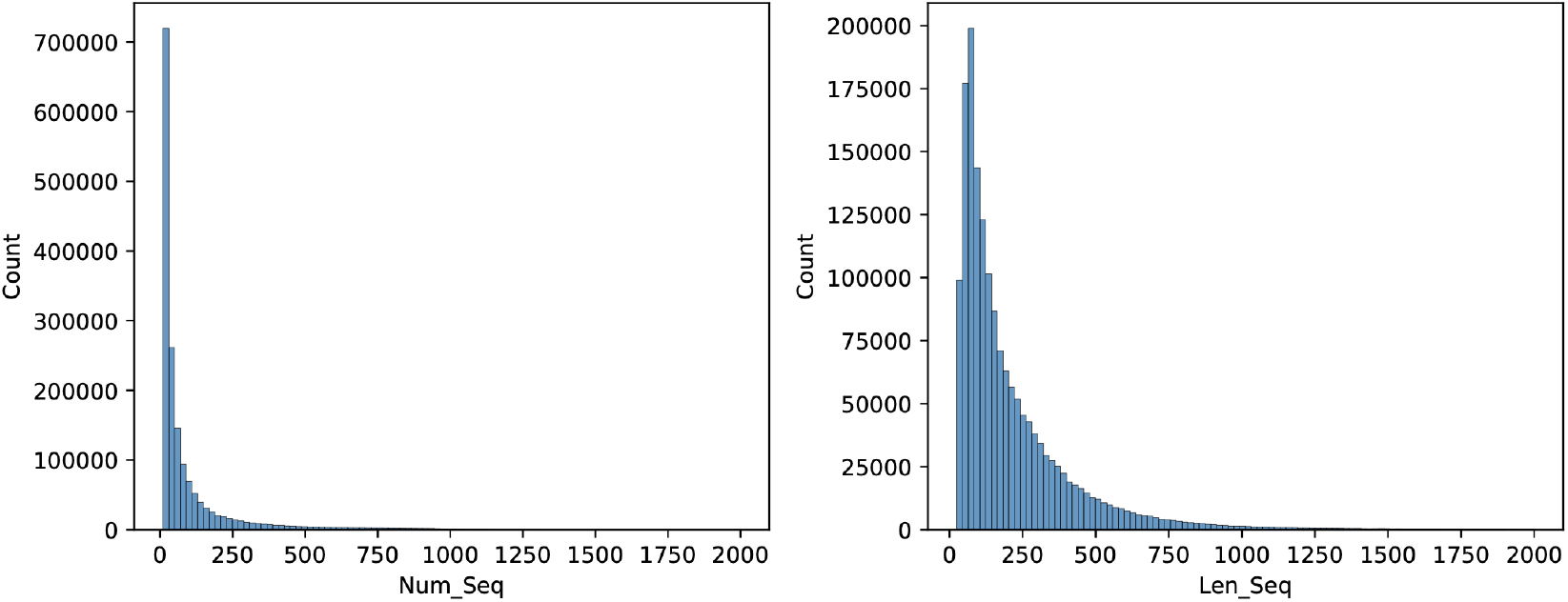
The length and depth distribution of the pre-training dataset.

### A.2 RFT

We fine-tune the base model using the pre-training cross-entropy loss on 𝒟_RFT_ with training only one epoch. Specifically, we adopt the same experimental settings as that used in the pre-training stage, except for the learning rate of 1.0×10^−5^ by default. Following the pre-training phase, the model undergoes a rejective fine-tuning process, which is more energy-efficient. This stage is executed on 8 x A800 GPUs for a single epoch for about two days.

#### RFT Dataset Filtering Threshold

When curating the RFT dataset, we first sample multiple MSA subsets for each protein structure, and select high-quality MSA subsets based on the following criteria: (1) the absolute structure prediction accuracy using the MSA subset, as measured by TM-score, should be larger than *θ*_1_, and (2) the relative improvement of the prediction accuracy after using the MSA subset, as compared to single sequence prediction, should be larger than *θ*_2_. We set *θ*_1_ = 0.9, and experiment with different *θ*_2_ values, as shown in table 4. The RFT model trained with dataset filtered by *θ*_2_ = 0.2 yields the best result, indicating that the relative information gain provided by MSA is a valuable indicator for curating high quality datasets for RFT. Moreover, *θ*_2_ = 0.5 results in a 20% decrease in dataset size, leading to inferior RFT model performance, highlighting the necessity of an intricate balance between data quality and data volumn.

**Table 4:** Performance comparison between different relative improvement threshold *θ*_2_ values.

| Threshold $\theta_2$ | 0 | 0.2 | 0.5 |
| --- | --- | --- | --- |
| MSAGPT+ RFT | 61.2 | **62.5** | 61.3 |

### A.3 RLAF

We fine-tune the RFT model using the DPO algorithm on 𝒟_DPO_ with training only one epoch. Specifically, we adopt the batch size of 1 with each MSA subset containing a maximum of 16,384 residues. We also use AdamW [49] with the learning rate of 1.0×10^−6^ by default. We linearly warmup the learning rate from 0 to 1.0 × 10^−6^ over the first 0.1% steps. This stage is also executed on 8 x A800 GPUs for a single epoch for about one day

#### RLAF Dataset

We conducted experiments with different data sources and filtering thresholds *θ*_3_—defined as the minimum relative improvement of the good case over the bad case in DPO data pairs—for the RLAF dataset, as detailed in Table 5. Utilizing only natural MSA subsets sampled from PDB, we found that higher *θ*_3_ values lead to improved model performance, suggesting a correlation between the disparity within data pairs and DPO effectiveness. Interestingly, the quality of MSA subsets generated by the RFT model surpasses that of natural MSA subsets at a *θ*_3_ of 0.2. However, the performance declines when natural MSAs are mixed with generated MSAs, compared to using a single data source during training. This indicates that maintaining distribution homogeneity is crucial for effective DPO alignment.

**Table 5:** Performance comparison between different data source and filtering threshold values.

| Data Source | nature(0.2) | nature(0.3) | generated(0.3) | nature(0.3)+generated(0.4) |
| --- | --- | --- | --- | --- |
| MSAGPT+ RLAF | 62.6 | **64.5** | 63.5 | 62.7 |

**Table 6:** The paired Student’s t-test between MSAGPT and other baselines on three benchmarks based on the TM-Score, where the p-value less than 0.05 indicates the result is said to be statistically significant.

| Model | CAMEO<br>(avg. Depth = 8.5) |  | CASP<br>(avg. Depth = 4.6) |  | PDB<br>(avg. Depth = 2.6) |  |
| --- | --- | --- | --- | --- | --- | --- |
|  | Zero-Shot | Few-Shot | Zero-Shot | Few-Shot | Zero-Shot | Few-Shot |
| AF2 MSA | 0.014 | 0.023 | 0.008 | 0.007 | 5e-7 | 8e-9 |
| MSA-Aug. | 0.023 | 0.014 | 0.044 | 0.015 | 6e-6 | 1e-7 |
| EvoGen | 0.038 | 0.027 | 0.067 | 0.016 | 1e-8 | 1e-9 |
| MSAGPT | - | - | - | - | - | - |

### A.4 Inference Efficiency

Generally, it’s vital to consider not just the immediate resource consumption during pre-training and post-alignment, but also the long-term utilization of these models. Once pre-trained, MSAGPT demonstrates significant efficiency, capable of generating protein sequences with up to 100,000 amino acids in under 8 hours. This efficiency underscores the model’s value, especially when amortized over its application lifespan and subsequent fine-tunings for specific tasks.

Regarding the scalability of the MSAGPT. We present the inference time with different total lengths (measured by protein sequence length multiply the number of generated sequences.), as shown in Figure 8.

**Figure 8:**
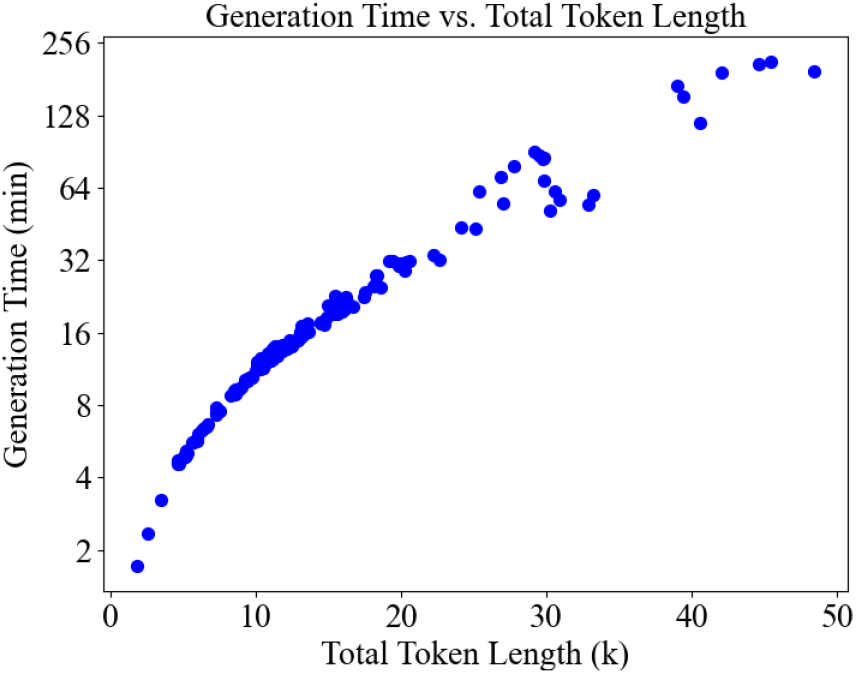
The correlation between total token length (the protein sequence length multiplied by the number of generated MSAs) and the inference time (minutes). In most cases (total token length < 20K), the generation time of MSAGPT is lower than the AF2 search pipeline requiring more than 30 minutes. The result shows MSAGPT can generate substantial sequence lengths within practical time, thus affirming its scalability and efficiency.

**Figure 9:**
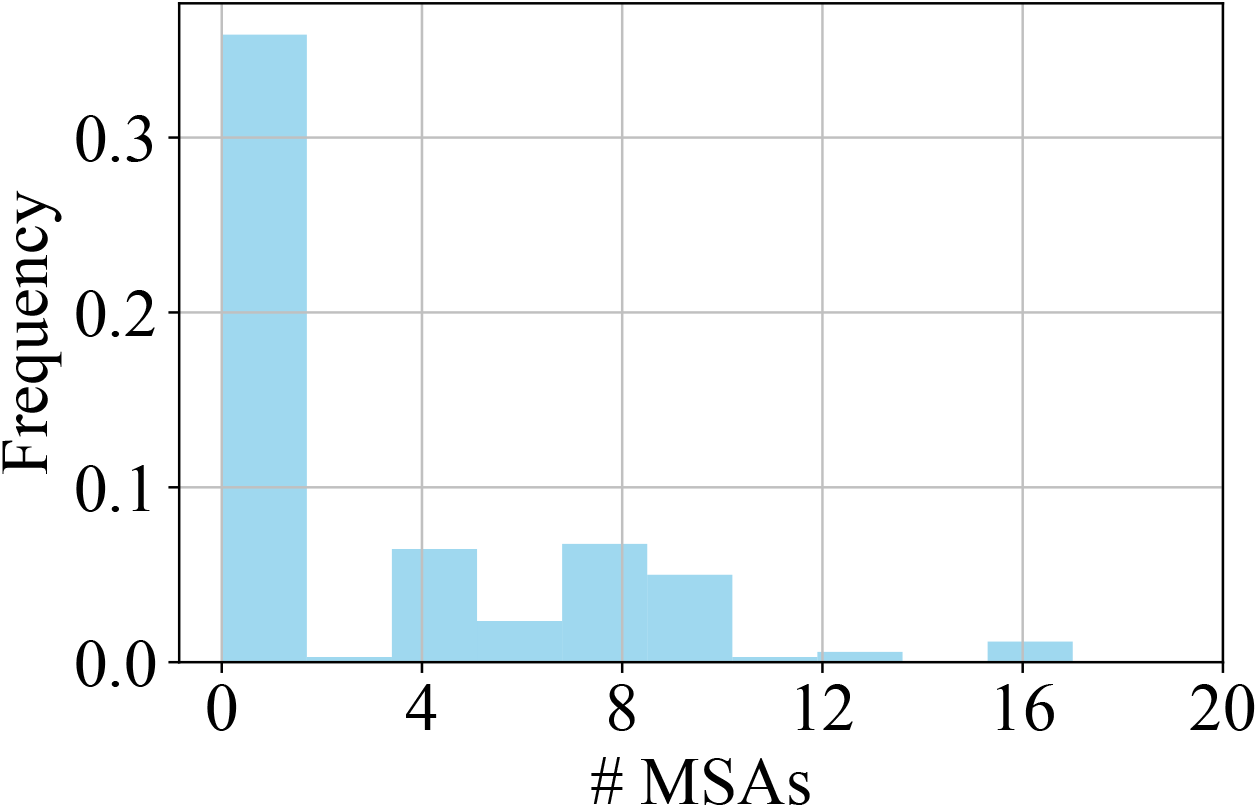
The distribution of MSA depth of benchmarked datasets.

The result showcases MSAGPT’s ability to generate substantial sequence lengths within practical time frames, thus affirming its scalability and efficiency.

## B Experimental Settings

### B.1 Evaluation details

We employ TM-Score, a widely-used metric for assessing the structural similarity between predicted structures and ground truth, and The predicted local distance difference test (pLDDT), a per-residue measure of local confidence. All metrics are scaled from 0 to 100, with higher scores indicating higher confidence and usually a more accurate prediction. where 1 indicates a perfect match between two structures. Each run across 3 independent runs. For each run, we adopt the different temperatures (T∈{0.8, 1.0}) along with different nucleus sampling factors (P ∈{0.8, 1.0}), experimenting with varying the number of generated MSAs in 8, 16, 32, and 64. The final performance is determined by first identifying the predicted structure with the highest accuracy across these different depths, and then averaging the results across the test set.

### B.2 Setup of Transferability of MSAGPT to Other Tasks

We utilized the MSA Transformer [17] as the backbone model with the task-specific heads. We finetune MSA transformer with the head with lr = 3*e* − 5 and batchsize = 16 on all experiments. All the task benchmarks are obtained following the pipeline in [11]. For each task, we sample 1000 protein sequences with the corresponding labels. Then we use MSAGPT-DPO to generate 32 virtual MSAs with the generation strategy T=0.8 and P=0.8. Both setups are trained briefly (for one epoch) for 5-fold cross-validation as shown in Table 7, and we report the average performance.

**Table 7:** The results of 5-fold cross-validation performance between with or without virtual MSA generated by MSAGPT on four protein-related tasks.

| Model | 1 | 2 | 3 | 4 | 5 | AVG |
| --- | --- | --- | --- | --- | --- | --- |
|  | Top (L/5) | ACC | ACC | ACC | ACC | - |
| w/o Virtual MSA (CtP) | 11.4 | 14.3 | 10.7 | 9.9 | 11.8 | 11.6 |
| **w/ Virtual MSA (CtP)** | 14.1 | 13.7 | 13.4 | 11.8 | 12.3 | **13.1** |
| w/o Virtual MSA (SsP) | 67.7 | 65.8 | 64.0 | 68.9 | 66.2 | 66.5 |
| **w/ Virtual MSA (SsP)** | 70.5 | 67.8 | 67.5 | 70.5 | 69.0 | **69.0** |
| w/o Virtual MSA (LocP) | 56.0 | 64.5 | 48.0 | 59.0 | 57.0 | **58.3** |
| **w/ Virtual MSA (LocP)** | 47.0 | 58.5 | 53.5 | 64.0 | 59.0 | 56.4 |
| w/o Virtual MSA (MIB) | 58.0 | 53.5 | 49.5 | 59.0 | 67.5 | 57.5 |
| **w/ Virtual MSA (MIB)** | 61.5 | 57.0 | 63.0 | 53.0 | 67.0 | **60.3** |

### B.3 Setup of Ablation Study

Experiments are conducted based on models with 150 million parameter size encompassing 30 layers, 20 attention heads, 640 embedding dimensions. These models are trained across approximately 30 billion tokens, amounting to around 40k training steps, maintaining consistency in hyper-parameter settings with MSAGPT, except for variations in the positional encoding mechanism. The efficacy of each variant is assessed through zero-shot MSA generation on the CASP test set.

## C Selection Strategy Details and pLDDT Evaluation

To evaluate the effectiveness of different selection strategies, we extracted 4, 8, 12, 16, 24, and 32 sequences from 48 zero-shot generated MSA for each method and computed the median TM-scores (Figure 4(b)) and pLDDT scores (Figure 10) across 33 test cases. The strategies are detailed below.

**Figure 10:**
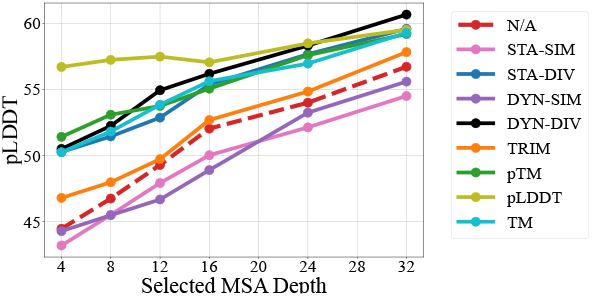
The pLDDT curves across different selection methods. Dashed red line represents using all generated sequences of a given depth. Solid lines represent selecting a subset of a given depth from 48 generated sequences with a specific strategy. The curves are smoothed using the Exponential Moving Average with alpha=0.3.

### Static Similarity / Static Diversity Strategy

We select the top-k generated MSA with the highest / lowest sequence identity to the query sequence. Sequence identity is determined by the percentage of identical residues between the two sequences.

### Dynamic Similarity / Static Diversity Strategy

Starting with the MSA most / least similar to the query sequence, we sequentially incorporate MSA into the selected set based on the highest or lowest average sequence identity with all sequences already included, until reaching a total of k MSA.

### Trimming Strategy

Suggested by EvoGen, this method filters out MSA with less than 50% coverage or sequence identity to the query sequence above 90% or below 20%. Subsequently, it iteratively selects the MSA with the closest sequence identity to the query and an average sequence identity below 90% with all the chosen MSA.

### pTM / pLDDT / TM Score Strategy

For each generated MSA, we remove the gaps and predict its structure using AF2. The structures are then ranked based on the pTM score (as reported by AF2), the pLDDT score (as reported by AF2), or the TM score compared to the query sequence’s ground truth structure (calculated by US Align), and the MSA corresponding to the top-k structures for each metric are selected accordingly.

## D Protein Structure Prediction Analysis compared with natural MSA

We present a detailed visual comparison of protein structures predicted by AlphaFold2 (AF2) utilizing MSA augmented by MSAGPT, against those predicted with natural MSA. This comparison, as depicted in Figure 12, highlights the remarkable capability of MSAGPTin enhancing the accuracy of structure predictions to levels that closely rival, and in some cases surpass, those based on naturally derived MSA.

We delve into a visualized analysis of protein structures predicted using AlphaFold2 (AF2) with MSA augmented by our proposed model (MSAGPT), alongside those augmented by EvoGen and MSA-Augmenter. This comparison, visualized in Figure 11, encompasses a spectrum of proteins, ranging from short sequences with relatively simple structures, like 7mnv_B, to long sequences with complex configurations, such as 7tdv_B. It includes proteins characterized by multiple beta sheets, exemplified by 7ywg_B, as well as those rich in alpha helices, such as 7tdv_B. Across these diverse cases, MSAGPT significantly surpasses both EvoGen and MSA-Augmenter, improving the TM score to a maximum of 0.9.

**Figure 11:**
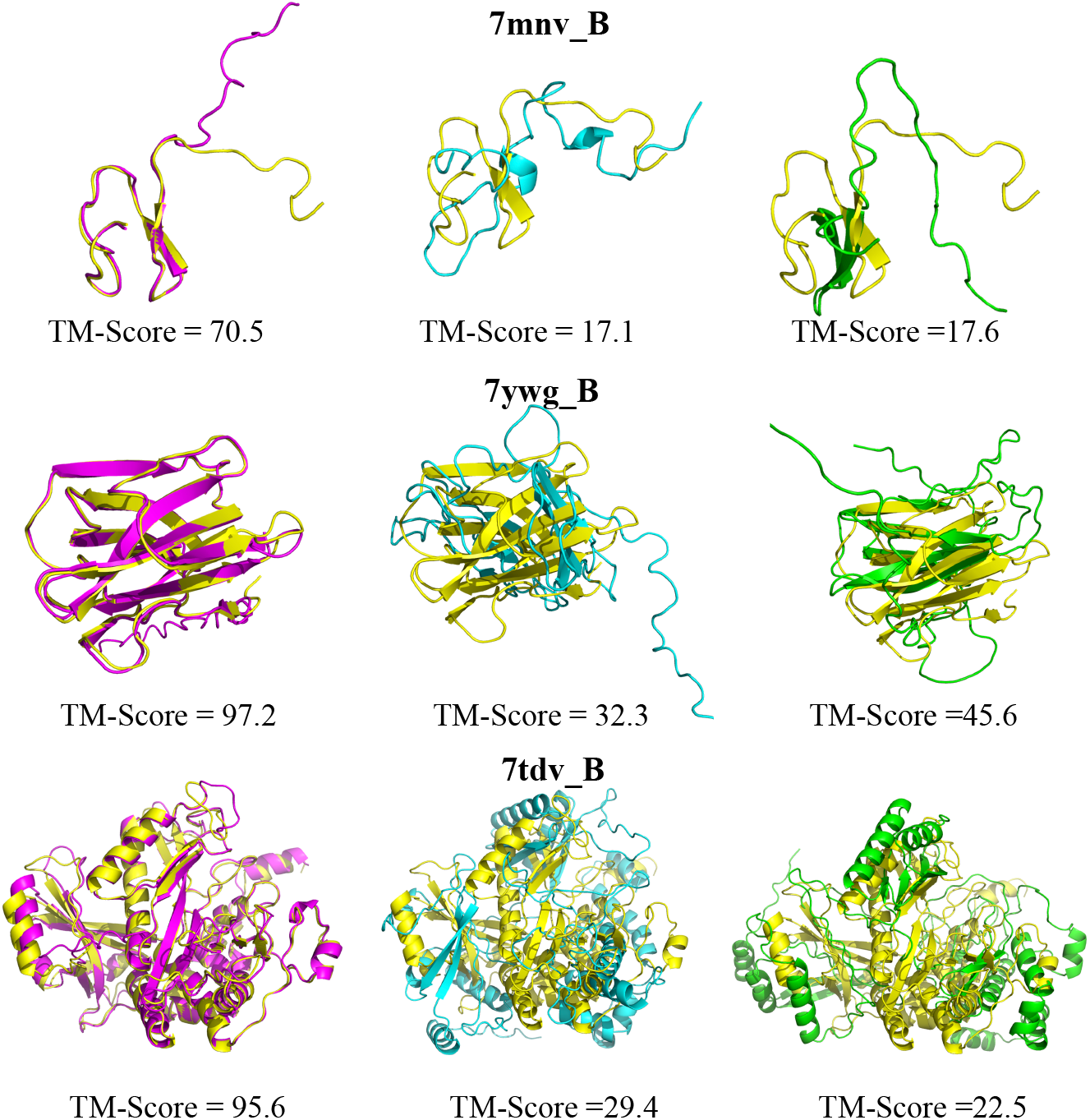
Visualization of improved structure prediction compared with baseline models. Yellow: Ground truth; Pink: Predictions based on MSA generated by MSAGPT; Blue: Predictions from MSA generated by EvoGen; Green: Predictions utilizing MSA generated by MSA-Augmenter.

**Figure 12:**
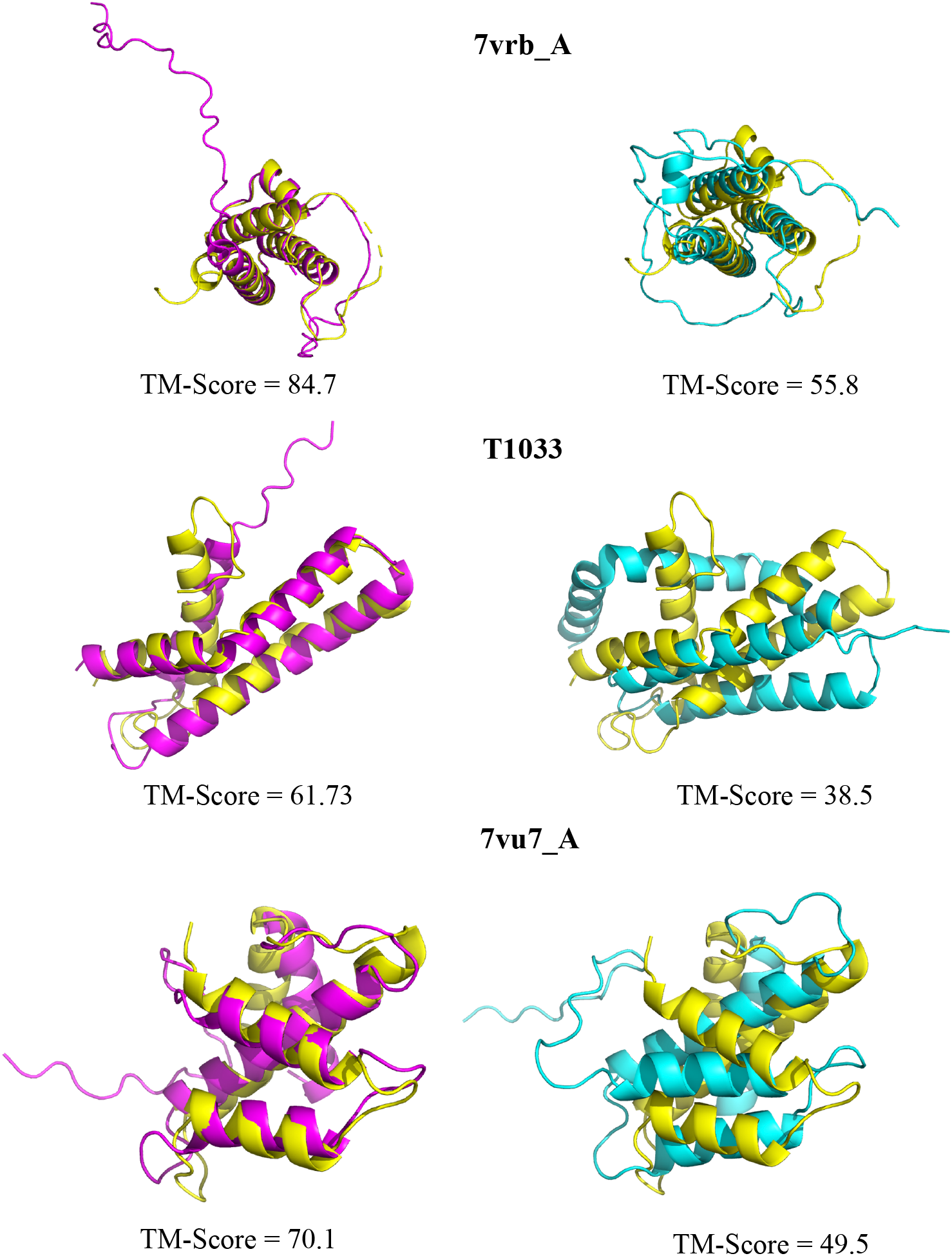
Visualization of improved structure prediction compared with nature MSA Yellow: Ground truth; Pink: Predictions based on MSA generated by MSAGPT; Blue: Predictions from MSA generated by natural MSA.

By detailed examination, we observe that while the MSA augmented by the baseline models assist AF2 in accurately predicting local structures and folds, they fall short in aligning the global composition and orientation with the ground truth structure, which is effectively addressed by MSA generated by MSAGPT. The local structures, which are generally more discernible from the spatial arrangements of adjacent amino acids, contrast with the global structures whose prediction relies heavily on comprehensively understanding the co-evolutionary information within MSA. These co-evolutionary patterns, indicating proximity in three-dimensional space through simultaneous mutations at multiple positions, are crucial for accurate global structure prediction. These findings underscore MSAGPT’s impressive capability to comprehensively capture and utilize co-evolutionary information, thereby significantly enhancing the accuracy of protein structure predictions. More visualization cases about the predictions based on MSA generated by MSAGPT and the predictions based on the natural MSA are illustrated in Appendix D.

## E Protein Structure Prediction Improvement after DPO

Figure 13 represents the comparison before and after the DPO training, depicting notable enhancements in structure prediction accuracy. Figure 14 and 15 provide an in-depth analysis of the generated MSA for each case. Specifically, residues 43, 53, 71-79, 105-111, 122, 132 and 157-166 in the MSA of 7wme_A, along with residues 22-27, 53, and 73 in the MSA of 7sxb_A, display distinct characteristics pre- and post-DPO training.

**Figure 13:**
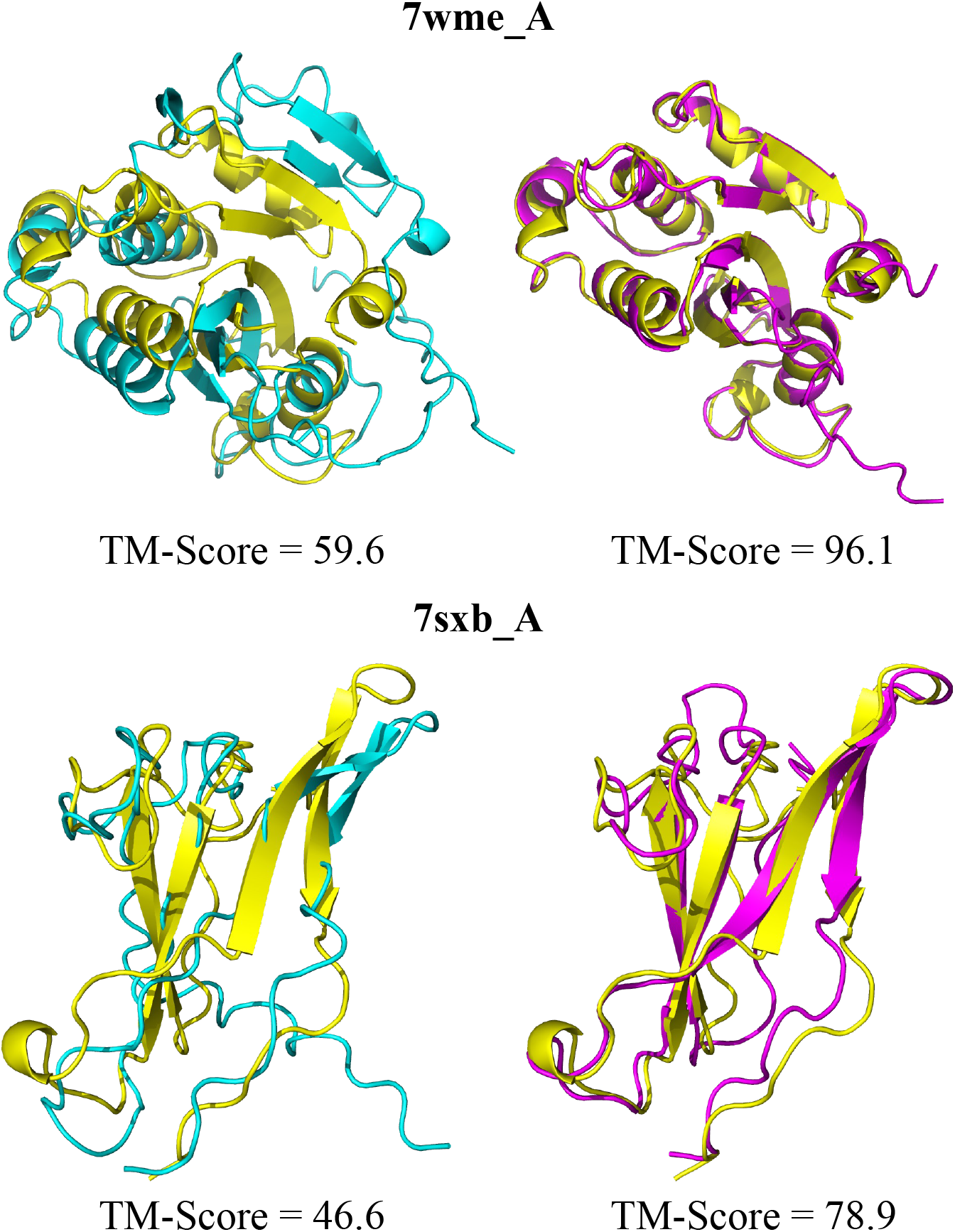
Visualization of improved structure prediction after DPO. Yellow: Ground truth; Blue: Predictions based on MSA generated by MSAGPT; Pink: Predictions based on MSA generated by MSAGPT-DPO.;

**Figure 14:**
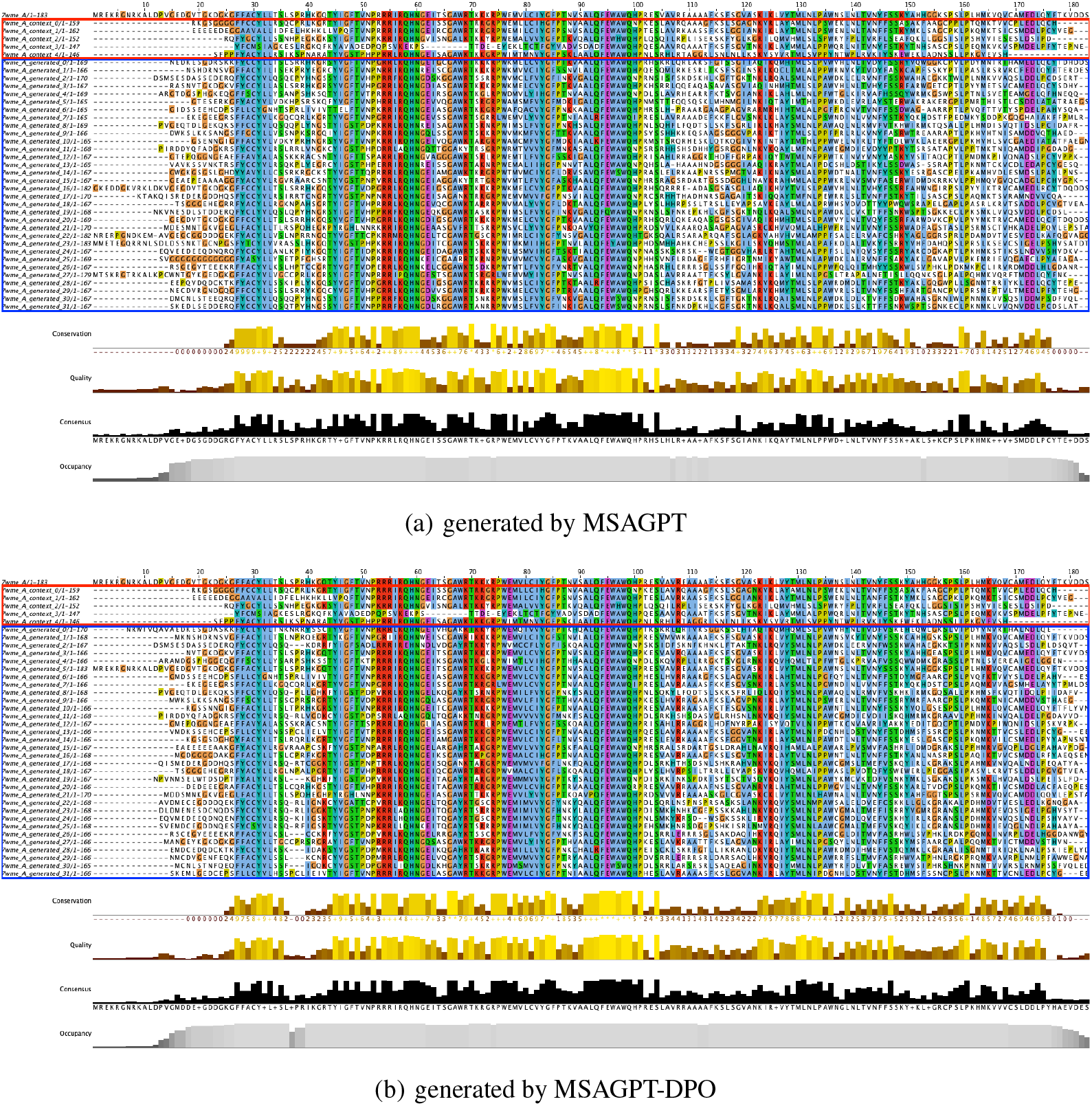
Residue Distribution of Generated MSA for 7wme_A. The red box indicates natural MSA used as prompts during generation. The blue box indicates generated MSA. Residues are colored using the clustal scheme by Jalview.

**Figure 15:**
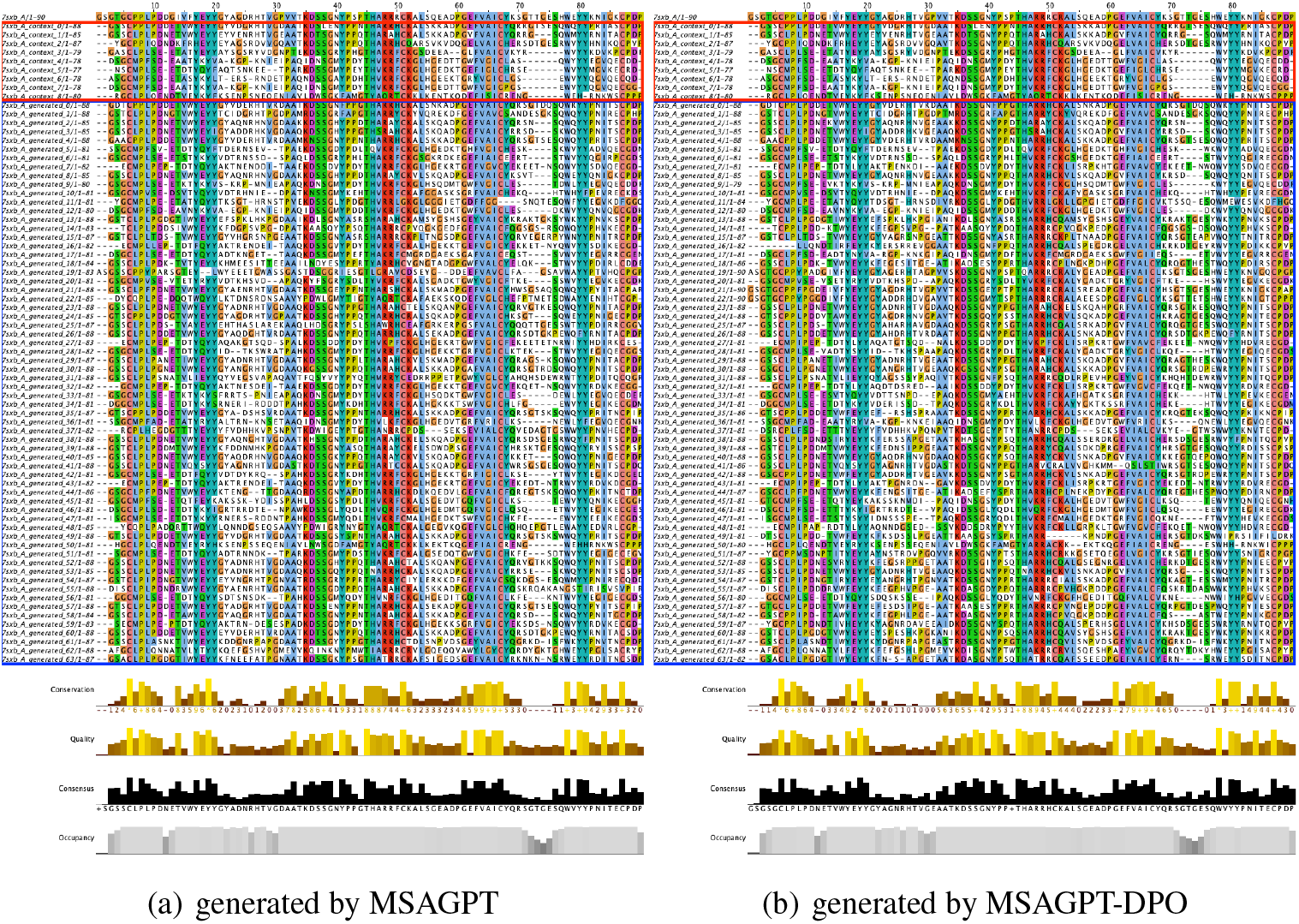
Residue Distribution of Generated MSA for 7sxb_A. The red box indicates natural MSA used as prompts during generation. The blue box indicates generated MSA. Residues are colored using the clustal scheme by Jalview.

## Footnotes

4 https://kexue.fm/archives/8397

